# Better Safe Than Sorry: Leg Amputations as a Prophylactic Wound Care Behaviour in Carpenter Ants

**DOI:** 10.1101/2025.06.29.662171

**Authors:** Seiji Fujimoto, Juan J. Lagos-Oviedo, Florian Seibel, Louis Puille, Ronja Hausmann, Eoin Corcoran, Thomas Schmitt, Erik T. Frank

## Abstract

Animals often sustain injuries, which are susceptible to lethal infections. In social insects, wound care behaviours have evolved to reduce these risks. But the limits of wound care behaviours remain unclear. Here we investigated the wound care behaviours of the ant *Camponotus maculatus*. Our findings show that amputation of legs infected with *Pseudomonas aeruginosa* significantly reduced mortality. However, nestmates do not differentiate between infected and sterile injuries, providing similar treatments regardless of infection. Even though we show that early amputation correlates with higher survival rates, nestmates amputate indiscriminately on legs with fresh or old wounds. Additionally, cuticular hydrocarbon profiles differed between ants with infected or sterile wounds only 24 hours post-injury, a timepoint when amputations are no longer effective. We propose that *C. maculatus* workers perform prophylactic amputations regardless of injury state or age. This is in sharp contrast to previous studies which showed clear capabilities to treat infected wounds differently in ants using antimicrobial compounds. This work therefore shows the limits of wound care behaviours in social insects, allowing us to better understand the evolutionary drivers of this unique behaviour.

## Introduction

In animals, injuries can occur in various contexts, such as hunting, foraging for food, or during inter- and intraspecific conflicts (Siva-Jothy et al. 2005; Mukherjee et al. 2013). Incurred wounds have a high likelihood of getting infected by opportunistic pathogens such as fungi and bacteria and pose a significant mortality risk in the animal kingdom (Hart 2011, Masson et al. 2024). To mitigate this risk, various animals have developed behavioural strategies to prevent infection such as grooming and licking wounds in mammals, to more sophisticated treatments such as the use of plant materials containing antiseptic compounds in some primate species (Kessler 2020). Injuries are not limited to vertebrates either, with social insects, like ants, having up to 20% of their workforce injured at some point in their life (Frank et al. 2017). To a higher extent than solitary animals, social insects such as honeybees, termites and ants face further vulnerabilities from wound infections as their densely populated nests favour the spread of pathogens, thereby threatening the entire colony’s wellbeing (Cremer et al. 2018, Schmid-Hempel 1998).

To counteract these increased disease risks, social immune responses encompass a set of collective defence adaptations to protect the colony against parasites and pathogens (Cremer et al. 2007). This can include behaviours like avoidance of infected nestmates, and modification of nest architecture to limit the spread of diseases within the colony after an infectious outbreak (Stroeymeyt et al. 2018, Leckie et al. 2024). Other strategies involve increased grooming among workers during pathogenic fungal outbreaks (Reber et al. 2011), or destructive disinfection of infected pupae (Pull et al. 2018). As well as protecting the whole colony from infectious threats, social immunity has evolved to prevent the loss of a significant part of the workforce which would seriously weaken the colony and threaten its survivability (Cremer 2018). One major cause of ant worker mortality is infection from injuries sustained during foraging (Frank et al. 2017, Frank et al. 2023).

Once thought to be a uniquely vertebrate behaviour, care toward injured individuals has been documented in various ant species. In the African termite-hunting ant *Megaponera analis*, workers injured during raids were carried back to their nests by their nestmates, where they received further wound care (Frank et al. 2017, Frank et al. 2018). If the wound was infected, nestmates further applied their antimicrobial metapleural gland secretions on the wounds of their injured, thus increasing the survivability of injured ants by up to 90% (Frank et al. 2023). In addition, *M. analis* workers can signal their injury state (whether infected or non-infected) to their nestmates by altering their cuticular hydrocarbon (CHC) profiles (Frank et al. 2023). A similar wound care behaviour was observed in the desert ant *Cataglyphis nodus* where treatments primarily appear to have a prophylactic role in preventing infections (Beydizada et al. 2024). In *Camponotus floridanus*, workers even performed amputations on injured legs of their nestmates, which significantly improved survival rates and stopped infections in injured workers (Frank et al. 2024).

The behaviour of amputating injured limbs to save a conspecific’s life, highlights the depth of ant social wound care systems and the necessity to investigate them further. Despite these promising findings, the limits of amputation’s effectiveness in ants remains unclear. For instance, do infected wounds receive more care than non-infected ones in *Camponotus*? How time-sensitive is the amputation process for it to successfully reduce mortality? Addressing these questions will provide deeper insights into the evolution and efficacy of wound care behaviours in social insects.

Here we describe the amputation behaviour and wound care treatment in the African ant *Camponotus maculatus* (Fabricius 1782). We investigate how wound infections with the gram-negative bacterium *Pseudomonas aeruginosa* affect wound care behaviours as well as the survival of injured workers. We further analyse the importance of the pathogen load and the timing of amputation for the survival of the injured as well as the role CHCs could play in communicating injury and infection.

## Materials and methods

### Study animal - laboratory maintenance conditions

The genus *Camponotus* is one of the most abundant and diverse ant genera worldwide, with 1,088 species described as of December 2024 (Bolton 2024). Among them, *Camponotus maculatus*, is found across Sub-Saharan Africa and the Arabian Peninsula (Borowiec 2014). This species nests in the ground, while its workers are frequently observed foraging in trees, where they tend to aphids (Taylor 1978).

Three colonies of *C. maculatus* were collected in Côte d’Ivoire (Ivory Coast) and established at the University of Würzburg in April 2019. All experiments were conducted in a climate chamber under conditions tailored to the natural habitat of *C. maculatus*. The temperature was maintained at 28°C with a relative humidity of approximately 80%. A 12-hour day-night cycle was simulated with light provided from 7:00 a.m. to 7:00 p.m. During the experiments, colonies were fed a honey-water nutrient solution, mealworms (*Tenebrio molitor*) and provided tap water *ad libitum*.

For behavioural experiments, sub-colonies of 100 ants were created from each of the three mother colonies. These sub-colonies included four major and 96 minor workers. Approximately half of the individuals were collected from within the nestbox, while the other half were collected from the foraging box. Each sub-colony was housed in a transparent plastic box (19 cm × 19 cm × 14 cm) with *ad libitum* food and water. The ants were given one week to acclimatize to the experimental setup. Twelve hours before the experiments, focal individuals were randomly chosen from each sub-colony and marked on their thorax with acrylic markers (Pebeo) for identification. For isolation experiments, each ant was kept in isolation chambers consisting of sterile petri dishes, filled with *ad libitum* water and honey-water.

### *Pseudomonas aeruginosa* culture

*Pseudomonas aeruginosa* is a gram-negative bacterium commonly associated with antibiotic resistance in humans (Kerr 2009), which also acts as an opportunistic pathogen ubiquitously present in the environment and able to cause deadly septicaemia in ants with infected injuries (Frank et al. 2023, Frank et al. 2024).

*Pseudomonas aeruginosa* cultures were prepared from a strain isolated from *M. analis* workers in Comoé National Park, northern Côte d’Ivoire, and cultured on tryptic-soy agar (Carl Roth GmbH + Co. KG, Karlsruhe) at 25**°**C for 18 hours. To ensure *P. aeruginosa* remained in the exponential growth phase, the bacterial culture was replated every 48 hours onto fresh nutrient medium. Infective solutions were prepared by collecting bacterial colonies from the edge of the culture and placed in 5 mL of sterile PBS. Optical density (OD) was measured using an Ultrospec 10 Cell Density Meter (Biochrom). Each solution was diluted until the desired OD concentration was reached.

### Wound care behavioural experiments

To compare the wound care behaviours towards infected and sterile-injured ants, we created 3 sub-colonies between November 8 and 21, 2022. Six previously marked ants were collected per sub-colony and all of them were wounded at the centre of the right hind femur using sterile scissors. Afterwards, half were exposed to a 10 μL pathogenic solution by immersing their injury in a solution containing approx. 10^5^ *P. aeruginosa* bacteria (0.01 OD) for 2 seconds. In the control group, the wound was exposed to sterile PBS for 2 seconds. All the workers were then returned to their respective sub-colonies for observations (N=9 per group).

The experiment was also repeated with a higher infection dose, infecting ants with a ten times more concentrated *P. aeruginosa* solution (0.1 OD, approx. 10^6^ *P. aeruginosa* bacteria). Observations followed the same procedure as above (N=9 per group).

Treated ants were observed in the nest for two weeks, recording the number of nearby ants, the treated worker’s condition (alive or dead), and overall colony mortality. Deceased ants were removed and inspected for limb amputation. Data on surviving ants and amputations was recorded every hour for the first 24 hours, every two hours for the remainder of the first week, and twice a day during the second week. Ant behaviour inside the nest was monitored using a Panasonic HC-X1000 4K video camera recorder mounted on top of the nest box. We analysed the first 6 hours of video recordings for each trial using VLC media player 3.0.16 Vetinari. We distinguished the following behaviours: (I) Wound care (injury is cleaned by another worker), (II) allogrooming (cleaning the injured worker), and (III) amputation (biting on the trochanter by a nestmate).

### Survival experiments

To confirm the role of amputation in reducing mortality rates associated with injury and wound infection, we compared the survival of injured ants with or without amputation and infection. A total of 80 workers (from 4 different colonies) were divided into 4 treatments. The first group labelled “Healthy,” served as a negative control and received no treatment (N=20). In the second group, “Sterile,” the right hind leg was severed at the centre of the femur using sterile scissors, and the wound was soaked in sterile PBS for 5 seconds (N=20). Workers in the “Infected” group were treated in the same manner as the “Sterile” group, but their wounds were soaked in a 0.01 OD *P. aeruginosa* solution for 2 seconds (N=20). The “Infected + Amputated” group underwent the same initial treatment as the “Infected” group, but after one hour, the leg was manually amputated at the trochanter using sterile scissors to simulate amputations performed by nestmates (N=20). All workers were kept in isolation chambers and provided *ad libitum* honey water and water. Isolation experiments were performed in parallel with a social experiment where 60 workers were collected from the same 4 colonies and received the same treatments as described above: (1) “Healthy”, (2) “Sterile” and (3) “Infected”. All the workers were then put back into their respective sub-colonies (N=20 per group). The isolation and colony survival experiments were repeated, using an 0.1 OD *P. aeruginosa* solution (N=20 per group). Workers were monitored once per hour for 6 days (144h) and the time of death was recorded.

To determine the time window during which amputations by nestmates remained effective in reducing infection mortality, we manually amputated the infected leg of injured ants kept in isolation at different timepoints. 10 workers (N=10) were collected per treatment group from four colonies. After manually injuring the femur and exposing the wound to a *P. aeruginosa* solution (0.1 OD), we amputated the infected leg at 0-, 6-, 12-, or 24-hours post-exposure. Mortality was quantified for the first 6 days post-exposure.

To investigate whether the duration ants carried an infected or sterile injury influenced the frequency of amputations by nestmates, we introduced injured nestmates at different timepoints after injury and/or infection into sub-colonies. Individual workers were collected from each of the 4 colonies and received one of three treatments: (1) “Healthy”, (2) “Sterile” or (3) “Infected”. Half of the workers were immediately returned to the nest after manipulation and the other half were held in isolation chambers with water and honey water provided *ad libitum* for 24 hours before being reintroduced to the nest (N=10 per group and time-point).

### Cuticular hydrocarbon analysis

A total of 70 workers were collected from one colony (N=70). Of these, 20 received a *P. aeruginosa* infected injury (OD 0.1), 20 received a sterile injury, and the remaining 30 did not receive any additional treatment as control (“healthy”). 10 individuals per group were frozen at -20°C, 2h and 24h after manipulation in addition to 10 healthy ants at 0h. Prior to freezing, the ants were kept alone in isolation in sterile petri dishes containing ad libitum water and honey water.

Cuticular hydrocarbons (CHC) were extracted from frozen workers by soaking their bodies in hexane for 10 minutes. The collected solution was then reduced to 100 μL for higher concentration before performing GC-MS analysis using a 6890 gas chromatograph (GC) with a 5975 Mass Selective (MS) Detector (Agilent Technologies, Waldbronn, Germany). The GC was equipped with a DB-5 capillary column (0.25 mm ID x 30 m; film thickness 0.25 μm, J & W Scientific, Folsom, CA, USA). A temperature program from 60°C to 300°C with 5°C/min and finally 10 min at 300°C was employed. Mass spectra were recorded in the EI mode with an ionization voltage of 70eV and a source temperature of 230°C. The area under the relevant peaks of the chromatograms were calculated using the software MassHunter Qualitative Analysis Navigator (v. B.08.00). Identification of the CHCs was accomplished by comparison of mass spectral data of commercially purchased alkane standards and diagnostic ions.

### Statistical analysis

For statistical analyses and graphical illustrations, we used the statistical software R v4.3.2 (R Core Team 2025) with the user interface RStudio 2023.06.0+421 (Rstudio Team 2020) and the R package ggplot2 v.3.3.5 (Wickham 2016).

To investigate behavioural differences between workers with infected and sterile wounds, we modelled wound care and allogrooming as binary response variables using Hierarchical Generalised Additive Models (HGAM) with a binomial family. A nested random effect with a smoother was included for each individual within a colony, along with a smoother interaction between time and wound condition. To identify time intervals with significant differences, we conducted post-hoc contrasts on the probability of these behaviours between infected and sterile ants. Given that wound care often occurs in short, intermittent bursts, we considered wound care to be present if it was observed for at least one-quarter of the observation interval (2.5 minutes). In contrast, allogrooming, which is more frequent during ant interactions, was considered present if it occurred for at least half of the observation interval (5.0 minutes). The conclusions remained robust regardless of variations in these criteria. Similarly, to determine when amputations were most likely to occur, we recorded the time at which amputated ants were first observed. Binary records per treatment were analysed using an HGAM model, as applied for allogrooming and wound care. Model assumptions were assessed using the *DHARMa* v.0.4.7 (Hartig 2024) and *mgcv* v.1.9.1 (Wood 2017) packages.

To determine whether ants with infected wounds undergo amputation sooner than those with sterile wounds, we used Generalised Linear Models (GLMs) to model the time to amputation as a function of wound condition with the package *lme4* v1.1.33 (Bates et al. 2015). Colony ID was included as a random effect to account for colony-level variation.

To evaluate the impact of amputation on mortality at low and high pathogen concentrations, we fitted separate Mixed Effects Cox Models to analyse censored survival data for each experiment. The models included treatment (healthy, sterile, infected, infected + amputated) and environment (colony, isolated) as additive effects, with Colony ID as a random effect to account for colony-level variation. Model fitting was performed using the *coxme* v.2.2.22 (Therneau 2024).

To assess at which time the amputations are no longer efficient in reducing mortality caused by infections, we fitted a Mixed Effects Cox Model for the censoring data on the survival of amputated ants at different times. Colony ID was included as a random effect. Furthermore, to validate that the mortality of workers was caused by the infections, we compared the survival of workers with an infected and sterile wound by cumulative Kaplan-Meier curves with the packages *Surv* v.3.5.5 and *survminer* v.0.5 (Kassambara et al. 2024). To identify differences in the mortality of specific groups, we performed pairwise comparisons on estimated marginal means. We controlled for false discovery rates on multiple comparisons by Tukey adjustments using the package *emmeans* v.1.11.1 (Lenth 2025).

CHC analyses were performed using the packages ggtext, gplots, GCalignR, ggdist, ggside and analyzeGC. Chromatographic data was aligned in R using the package GcalignR v.1.0.7 (Ottensmann et al. 2018). Subsequently, all compounds with a relative abundance below 0.1% or less than 50% occurrence within all groups were excluded. The final compounds were then identified, and a retention index was calculated based on Carlson et al. (1998).

To visualise dissimilarities between wound types over time, we calculated the Bray-Curtis dissimilarities indices for the relative abundance data between CHC samples. We further visualise these dissimilarities by performing non-metric multidimensional scaling (NMDS). To test whether ants sustaining infected wounds show a distinct CHC profile, we assessed differences between groups over time by performing a multivariate analysis of variance (PERMANOVA) with 9999 permutations. These analyses were performed with the package vegan v.2.6-10 (Oksanen et al. 2025).

## RESULTS

### Sterile and infected injuries are treated equally

To investigate whether ants care more for infected than sterile wounds, we estimated the probability of receiving wound care and allogrooming in the nest during the first six hours after injury (figure 1a). Even though isolated ants with an infected wound had a 90% mortality after 18 days compared to a 40% mortality in isolated ants with a sterile wound (figure S1; Hazard ratio: z = -1.99, p = 0.05), we observed that inside the nest ants were equally likely to receive wound care and allogrooming regardless of wound condition (figure 1a,b, figure S2; wound care HGAM: z = 0.53, p = 0.60; allogrooming HGAM: z = -0.35, p = 0.73). For both wound conditions, wound care and allogrooming decreased over the first 6 hours after injury (figure 1a,b). Both wound conditions also showed a similar rate and timing of amputation (figure 1c, proportion of amputations, Chi^2^ test: χ^2^ < 0.001, df = 1, p = 1; amputation timing, GLM: *F* = 0.15, df = 1, p = 0.72). When wounded ants were immediately returned to the sub-colony, infected wounds were amputated in 100% of the cases (N= 10), and sterile wounds in 80% of the cases (Chi^2^ test: χ^2^ = 0.56, df = 1, p = 0.46, N= 10). A similar amputation pattern was observed for ants with 24 hours old wounds when reintroduced to their sub-colonies (infected wounds 100%, sterile wounds 90%; Chi^2^ test: χ^2^ <0.001, df = 1, p = 1; N=10 per group). Wound age thus did not affect amputation rates (Chi^2^ test: χ^2^ = 3.96, df = 3, p = 0.27).

**Figure 1.**
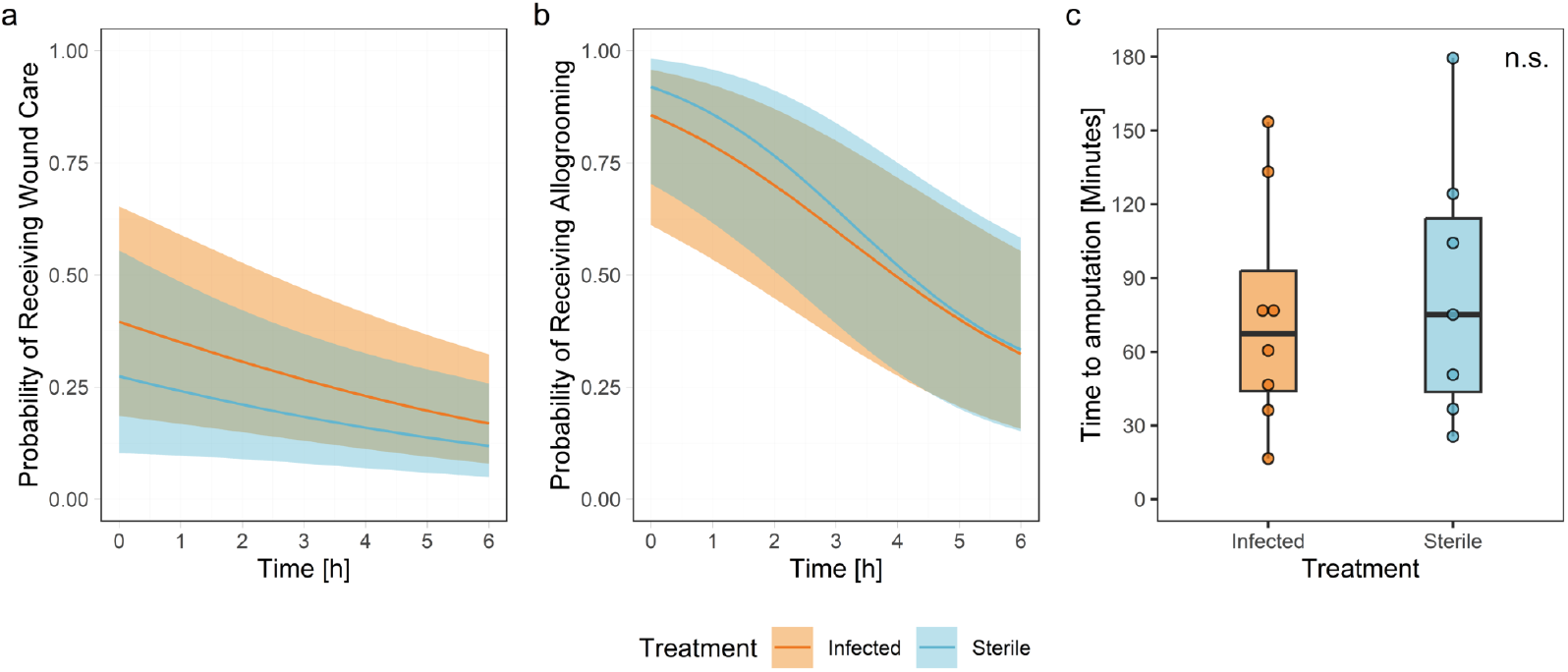
Workers received the same level of wound care, allogrooming and amputation regardless of wound condition. **(a)** Probability of an ant receiving wound care for infected and sterile wounds. The interaction between time and treatment (smooth term) was significant for infected (p < 0.001, df = 2.26, χ^2^ = 27.79) and sterile wounds (p < 0.01, df = 3.24, χ^2^ = 17.64). The Hierarchical Generalised (binomial) Additive Model (HGAM) explained 29% of deviance, with a coefficient of determination *R*^2^ = 0.17 over 667 observations. N = 9 per treatment. The curve represents the predicted probability for the behaviour by the HGAM, with the shaded area representing the 95% confidence intervals. **(b)** Probability of an ant receiving allogrooming for infected and sterile wounds. The interaction between time and treatment (smooth term) was significant for infected (p < 0.001, df = 4.71, χ^2^ = 55.51) and sterile wounds (p < 0.001, df = 6.05, χ^2^ = 62.70). HGAM explained 39% of deviance, with a coefficient of determination *R*^2^ = 0.40 over 667 observations. N = 9 per treatment. **(c)** Timepoint when ants with infected or sterile wounds received amputations in the colony. Regardless of the injury condition, workers receive amputations at similar times (GLM: *F* = 0.15, df = 1, p = 0.72). Boxplots depict the first to third quartile of the interquartile range, horizontal lines within boxes are medians, and the whiskers are 1.5 interquartile range. Within the first six hours of observation, 8 out of 9 workers received amputations in the infected group, and seven out of nine in the sterile group.

### Amputation efficacy decreases with a higher pathogen load

To assess how amputations help mitigate infections at different pathogen loads, we compared the survival of workers who were manually amputated after receiving a high or low dose of pathogens with those who did not undergo amputation. Our results showed that amputations after 1 hour reduced mortality by 66% in ants exposed to a low pathogen concentration compared to non-amputated infected ants (figure 2a, figure S3a; Hazard ratio: z = 2.33, p < 0.05). Similarly, amputations reduced mortality by 51% in ants exposed to high pathogen concentrations compared to non-amputated infected ants (figure 2a, figure S3a; Hazard ratio: z = 2.85, p < 0.05). While ants exposed to a low pathogen concentration after amputation had a similar mortality curve compared to ants carrying a sterile wound (figure 2a, figure S3a; Hazard ratio: z = -0.95, p = 0.34), ants that were exposed to a high pathogen concentration after amputation were still more likely to die than ants with a sterile wound (figure 2a, figure S3a; Hazard ratio: z = -2.73, p < 0.05).

**Figure 2.**
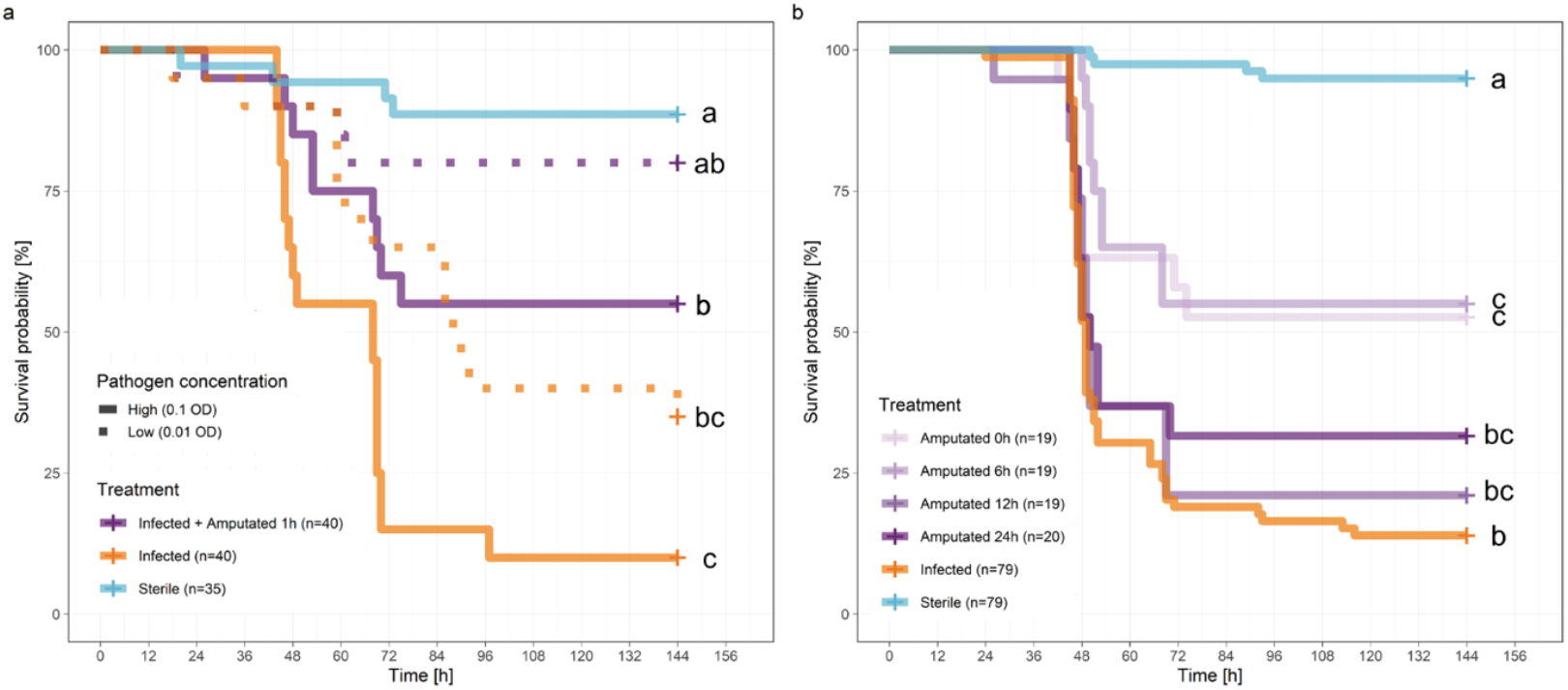
Survival probability with and without amputations. **(a)** Kaplan-Meier cumulative survival curves illustrate the effectiveness of amputations in mitigating varying levels of pathogen infections in isolated ants. Data were collected from four colonies: N_sterile_= 35, N_other groups_= 20. Statistical analyses and model comparisons were conducted using a Mixed Effects Cox Model (table S1, S2) **(b)** Kaplan-Meier cumulative survival curves of ants that received an infected injury (OD=0.1) followed by an amputation at different times. N_sterile_= 79, N_infected_= 79, N_amputated 24h_= 19, N_amputated 12h_= 19, N_amputated 6h_= 20, N_amputated 0h_= 19. Statistical analyses and model comparisons were conducted using a Mixed Effects Cox Model (table S3). Different letters depict statistical differences between groups for the estimated marginal means for p < 0.05.

Both under low and high pathogen concentration, the survival of infected ants kept inside the colony was similar to that of individuals who were experimentally amputated after 1 hour in isolation (figure S4a,b, table S1, Hazard ratio low concentration, z = -1.27, p = 0.21; Hazard ratio high concentration: z = -0.75, p = 0.45). Nestmates amputated the injured leg within the first 2 hours after introduction in both cases in a similar frequency (Low concentration: 79% amputated, Chi^2^ test: χ^2^ = 0.20, df = 1, p = 0.66, N= 19; High concentration: 80% amputated, Chi^2^ test: χ^2^ = 1.82, df = 1, p = 0.18, N= 20). Notably, infected ants that received an amputation by nestmates in the nest were much more likely to survive (88% after 60 hours, N= 15) than ants that were not amputated (25% after 60 hours, N= 4) at both high and low concentrations (figure S4c,d, table S2).

### Amputations lose effectiveness over time

To identify when amputations are no longer effective at reducing infection mortality, we experimentally amputated the legs of infected ants at 0, 6, 12, and 24 hours after pathogen exposure (OD=0.1). Ants that received amputations at the moment of infection or 6 hours later had a 48% reduction in mortality compared to ants that were not amputated (figure 2b, figure S3b; Hazard ratio Infected-Amputated-0h: z = -2.79, p < 0.01; Hazard ratio Infected-Amputated-6h: z = -3.43, p < 0.001). When amputations were delayed 12- or 24-hours mortality did not differ from infected ants that did not receive amputation (figure 2b, figure S3b; Hazard ratio Infected-Amputated-12h: z = -0.73, p = 0.47; Hazard ratio Infected-Amputated-24h: z = -1.29, p = 0.20). These results indicate that the timing of amputation is critical for improving the survival chances of infected ants.

### Changes in CHC profiles following wounding and infection

To determine whether workers’ cuticular hydrocarbon (CHC) profiles change after wounding and infection, we collected CHCs from ants with infected and sterile wounds, zero, 2- and 24-hours post-injury. We identified 39 cuticular hydrocarbons, predominantly alkenes (66.0%), followed by alkanes (33.4%), and a small amount of methyl-branched alkanes (1.1%; table S4) and alkadienes (0.1%; table S4). Our analysis revealed that CHC profiles changed depending on wound state and time (figure 3, table S5, PERMANOVA: F = 3.99, R^2^ = 0.10, p < 0.01). These differences were primarily observed 24 hours after injury between ants with infected or sterile wounds (table S6, pairwise comparison: R^2^ = 0.43, p = 0.02). However, neither wounded group differed significantly from healthy control ants after 24 hours (table S6, sterile-healthy: R^2^ = 0.15, p = 0.12; infected-healthy: R^2^ = 0.16, p = 0.12). At two hours, when most of the amputations typically had already occurred in the colony, we detected no differences between the CHC profiles of ants with infected or sterile wounds (table S6, pairwise comparison: R^2^ = 0.02, p = 0.80), nor did wounded ants differ in their profile compared to healthy ants (table S6, sterile-healthy: R^2^ = 0.02, p = 0.80; infected-healthy: R^2^ = 0.04, p = 0.49).

**Figure 3.**
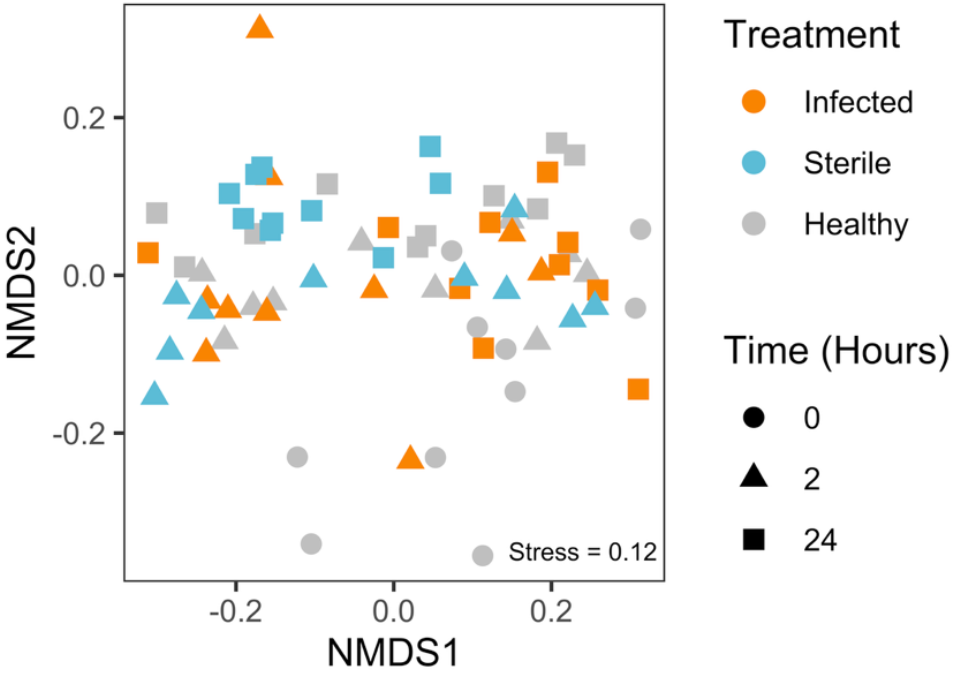
Changes of CHC profiles of infected and sterile ants over time. Bi-dimensional Non-metric Multidimensional Scaling (NMDS) analysis on a dissimilarity matrix of the CHC profiles of ants with infected (orange) or sterile (blue) wounds at different times after treatment compared to healthy ants (grey). N= 10 per time-point and treatment.

## DISCUSSION

Our study reveals that wound care in *C. maculatus*, including amputation behaviour, significantly reduces mortality in injured ants and functions as a prophylactic rather than a therapeutic measure. Workers do not discriminate between sterile and infected wounds or between fresh and old injuries, instead treating all wounds similarly through amputations. We found that high pathogen loads correlate with increased mortality and that amputation is most effective within a narrow time window post-injury. Finally, CHC profiles showed little change following injury, thereby corroborating our behavioural results that workers cannot detect their nestmates’ infection status and instead treat all injuries prophylactically.

Similar to *C. maculatus*, workers of *C. floridanus* showed comparable amputation rates between infected and sterile femur injuries (Frank et al. 2024). However, wound location (tibia or femur wound) induced different wound care responses in *C. floridanus* (Frank et al. 2024). Other ant species seem to be able to differentiate infected from sterile injuries. Workers of *Cataglyphis nodus* with infected wounds received no further care from their nestmates and were even expelled from the nest, while workers with sterile injuries received continuous wound care by their nestmates (Beydizada et al. 2024). In *M. analis*, nestmates treated workers with infected injuries with additional care by using antimicrobial secretions from their metapleural gland to actively combat the infection (Frank et al. 2023).

We found that amputations significantly improved survival following both low and high pathogen exposures, though its effectiveness was lower at high pathogen loads. Ants amputated after low-dose infections (0.01 OD) survived at rates comparable to those with sterile wounds, whereas those amputated after high-dose infections (0.1 OD) still experienced an elevated mortality. This suggests that while amputations can mitigate septicaemia, its success is constrained by the severity of the initial pathogen load. However, infections with a single species pathogen load as high as 10^6^ bacteria in 10μL are unlikely to occur in the wild. An equivalent 10μl droplet of surface soil likely contains between 10^5^ to 10^6^ bacteria of a diverse community that likely also inhibit each other when exposed on a wound (Raju et al. 2017). We can thus assume that wound care behaviours of *C. maculatus* are effective at handling injuries occurring under natural conditions where extremely high pathogen loads are unlikely to occur.

Workers also do not differentiate between fresh and old wounds, amputating at similar rates immediately or 24 hours after injury. However, our data shows that the longer injured workers are left alone without treatment, the higher the mortality. There appears to be a critical window between 6 and 12 hours, beyond which amputation is no longer effective, with survivability dropping below 25%, a rate comparable to workers which did not receive amputation. While isolation in ants and insects has been associated with stress and a weakening of resistance to pathogens (Kohlmeier et al. 2016), one significant factor in *C. maculatus* seems to be the spread of *P. aeruginosa* pathogens from the leg into the rest of the body within 6 to 12 hours. As our data shows that survival depends on timely treatment, waiting for infection signals to become detectable would thus be detrimental. Instead, assuming the likely worst case (an infected wound) and treating all injuries in the same way, amputation, is an effective strategy to maximise survival of injured workers. Furthermore, since amputations seem to have no added cost for the survival of injured ants, there are no clear disadvantages of performing amputations on injured individuals indiscriminately. The selection pressure to assess wound state or age to maximise worker survival in *C. maculatus* is thus likely extremely low. Finally, under natural conditions, the occurrence of injuries older than 6 hours before encountering nestmates is highly unlikely.

Workers of *C. maculatus* likely cannot detect infections through changes in the CHC profiles of injured nestmates on time for amputations, since no significant changes were observed between the CHC profiles 2 hours after injury and infection. Workers of *M. analis*, which adapt their treatment depending on the state of the wound, show changes in their CHC profile 2 hours after wounding and clearly detectable differences between sterile and infected wounds after 11 hours (Frank et al. 2023). These changes in *M. analis* mostly occur via differences in abundance of alkadienes, which are known to be involved in communication and signalling in ants in general (Sprenger 2020). Interestingly, in *C. maculatus* CHC profiles after 24 hours differed between workers with infected and sterile injuries, suggesting that once the immune system is fully activated in response to an infection, diseased individuals could be identified within the colony, potentially to isolate diseased individuals or allow for further care (Cremer et al. 2018, Stroeymeyt et al. 2018).

In light of these results, we conclude that amputation behaviour serves a prophylactic role rather than an active therapeutic function. This contrasts with species like *M. analis*, which invests more into treating infected wounds, and *Cataglyphis nodus*, where workers receive treatment only for recoverable injuries while infected individuals are left untreated (Frank et al. 2023; Beydizada et al. 2024). Given that *C. maculatus* lacks metapleural glands and likely cannot actively treat infections once they spread, amputation may function as an effective prophylactic strategy to mitigate mortality and workforce loss. The wound care behaviours observed in *C. maculatus*, and likely shared by its close relative *C. floridanus* (Frank et al. 2024), represents one of the first clearly documented cases in insects where amputations are consistently applied as a prophylactic strategy against wound infections, with experimentally tested limits and rationale for its use. The selectiveness in use of metapleural gland secretions in *M. analis* during wound care suggests a cost to antimicrobial production and application, possibly explaining why it can differentiate wound states. Since opportunistic pathogens such as *P. aeruginosa* are very common in the environment and injuries are very likely to become infected in the wild (Frank et al. 2023), workers in *C. maculatus* and *C. floridanus* should respond with indiscriminate prophylactic amputations, which come at no significant cost. However, if we assume that amputations carry little cost and are highly effective, and that most ant species are exposed to the same risk of wound infections, it remains unclear why amputation is present as a wound care strategy in some ant species (*C. maculatus* and *C. floridanus*) but not in others (*M. analis* and *Cataglyphis nodus*). We therefore hypothesize that some species such as *M. analis* and others in the Ponerinae subfamily may be unable to perform amputations due to their approximately ten times thicker cuticle (Peeters et al. 2017), which likely evolved to protect them from more severe injuries while hunting pugnacious prey. Thus, morphological features, as well as general ecology, may play a role in shaping the evolution and diversity of wound care strategies in ants.

## Competing interests

The authors declare no competing interests.

## Acknowledgements

We thank the Comoé National Park Research Station for the use of their facilities and the park management of Office Ivoirien des Parcs et Réserves (OIPR) for facilitating the collection of the ants. We thank David Kouassi and Abou Ouattara for their help collecting the ant queens.

## Funding

This work was supported by the Emmy Noether Programme of the German Research Foundation [grant number: 511474012].

## Notes

### Competing Interest Statement

The authors have declared no competing interest.

