## Supplementary Figures and Tables for "Better Safe Than Sorry: Leg Amputations as a Prophylactic Wound Care Behaviour in Carpenter Ants"

Affiliations:
[1] Department of Animal Ecology and Tropical Biology, University of Würzburg, Würzburg, Würzburg, Germany
[2] Department of General Systems Studies, Graduate School of Arts and Sciences, The University of Tokyo, Tokyo, Japan
[3] Department of Conservation Biology and Forest Ecology, University of Würzburg, Würzburg, Germany

*Shared first authorship

[+]Corresponding authors:
Juan J. Lagos-Oviedo:
Erik T. Frank:

### Supplementary Figures

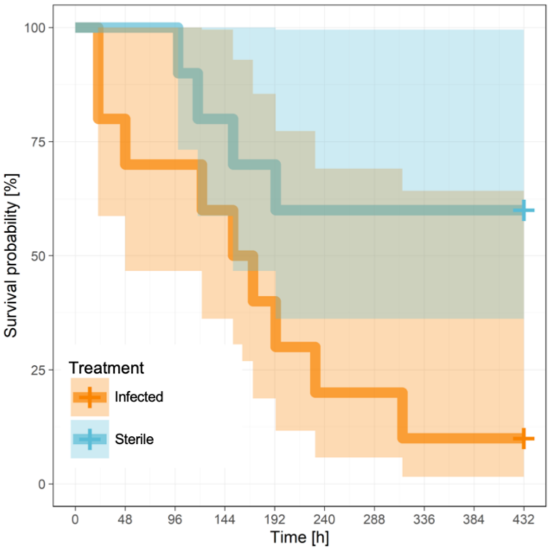


**Figure S1. Kaplan-Meier survival curves for infected and sterile injured ants for the behavioural observations in the nest.** Ants with sterile injury were significantly less likely to die than infected ants (Hazard ratio sterile: $z = -1.99$, $p = 0.05$). Shading indicates 95% confidence intervals. ($N=10$ per group).

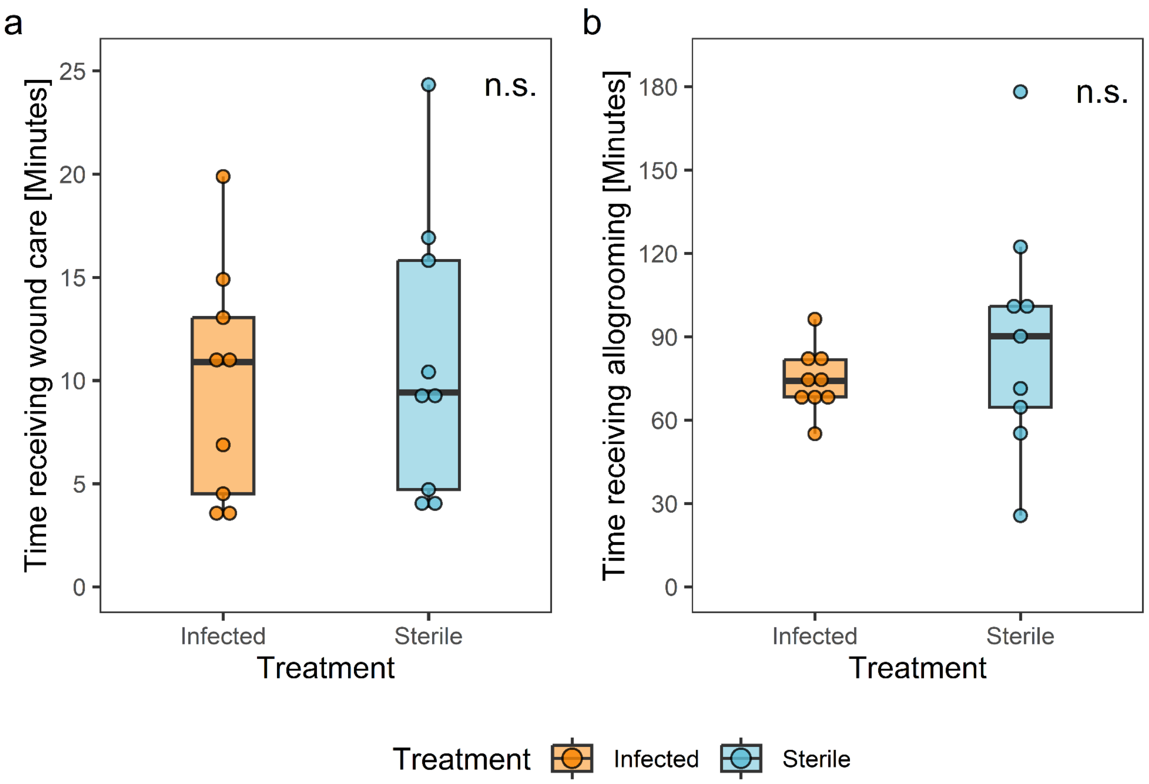


**Figure S2. Duration of wound care and allogrooming received by ants with infected versus sterile wounds**. **(a)** Injured workers received wound care for a similar amount of time regardless of the injury condition (GLM: $F = 0.15$, df = 1, p = 0.70). **(b)** Injured workers received similar amounts of allogrooming regardless of the injury condition (GLM: $F = 1.06$, df = 1, p = 0.32). Boxplots depict the first to third quartile of the interquartile range, horizontal lines within boxes are medians, and the whiskers are 1.5 interquartile range. n.s.= not significant (p>0.05); N=9 per group.

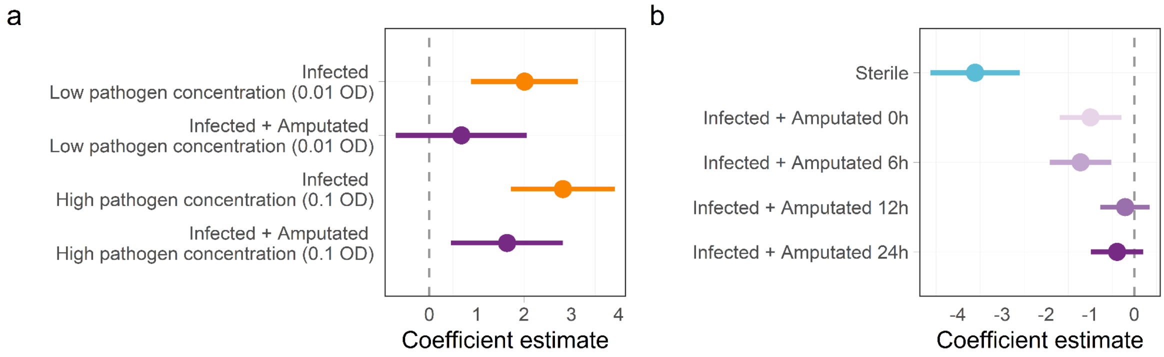


**Figure S3. Estimated hazard ratio coefficients for the survival with and without amputations. (a)** Effect of pathogen concentration on amputation efficacy. The forest plot shows the relative risk ratios of amputations in mitigating varying levels of pathogen infections in isolated ants (table S1, S2). Coefficients were compared to ants with a sterile wound as the reference group (dotted line). **(b)** Time limits for successful amputations. The forest plot shows the relative risk ratios of amputations at different times after the infection of a wound (table S3). Coefficients were compared to ants with an infected wound as the reference group (dotted line). Dots represent the estimated hazard ratio with a 95% confidence interval.

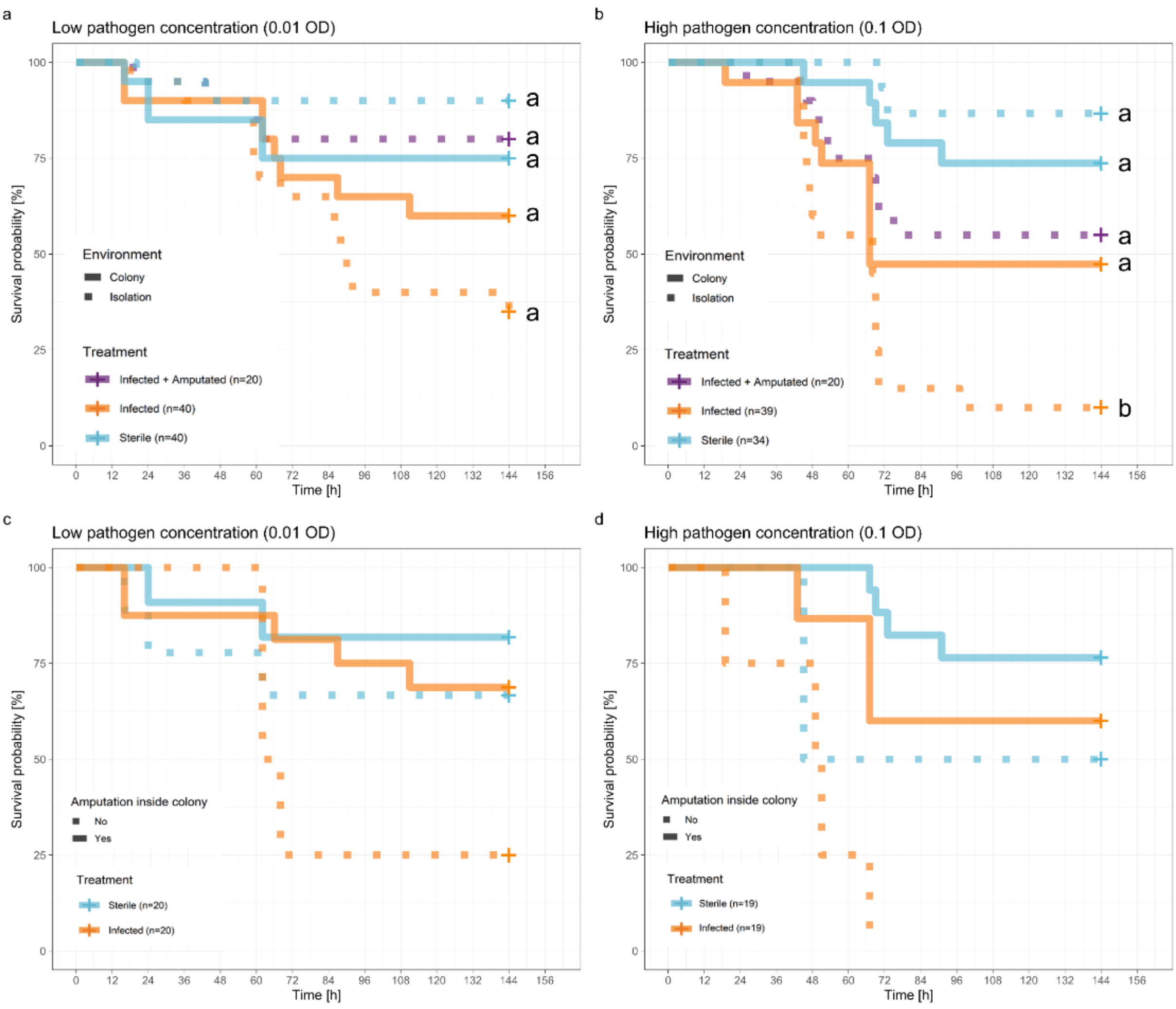


**Figure S4. Worker survival at different pathogen concentrations inside sub-colonies.** Kaplan-Meier cumulative survival curves illustrate the effectiveness of amputations in mitigating varying levels of pathogen concentrations. **(a)** Low pathogen concentration (0.01 OD). Sample size $N_{colony + infected}$= 20, $N_{colony + sterile}$= 20, $N_{isolated + infected}$= 20, $N_{isolated + sterile}$= 20. **(b)** High pathogen concentration (0.1 OD). Sample size $N_{colony + infected}$= 19, $N_{colony + sterile}$= 19, $N_{isolated + infected}$= 20, $N_{isolated + sterile}$= 15. Survival for ants that did or did not receive amputation inside the sub-colony at **(c)** low pathogen concentration (0.01 OD). Note that the frequency of non-amputations was low: $N_{infected + non\text{-}amputated}$ = 4, $N_{infected + amputated}$=16, $N_{sterile + non\text{-}amputated}$=9, $N_{sterile + amputated}$=11. **(d)** Survival for ants that did or did not receive amputation inside the sub-colony at high pathogen concentration (0.1 OD). Note that the frequency of non-amputations was low. $N_{infected + non\text{-}amputated}$ = 4, $N_{infected + amputated}$=15, $N_{sterile + non\text{-}amputated}$=2, $N_{sterile + amputated}$=17. Statistical analyses and model comparisons were conducted using a Mixed Effects Cox Model. Different letters indicate statistically significant differences in estimated marginal means at $p < 0.05$.

### Supplementary Tables

**Table S1.** Statistical differences in ant survival across different pathogen concentrations. Ants infected within the colony were used as the reference level for both comparisons. We fitted a Mixed Effect Cox proportional hazards model by adding the colony as an intercept random effect.

| Comparison | estimate | std.error | statistic | p-value | conf.low | conf.high |
|---|---|---|---|---|---|---|
| *Low pathogen concentration* (0.01 OD) | | | | | | |
| Amputated-Isolated | 0.460 | 0.613 | -1.267 | 0.205 | 0.138 | 1.530 |
| Infected-Isolated | 1.561 | 0.455 | 0.978 | 0.328 | 0.640 | 3.807 |
| Sterile-Colony | 0.620 | 0.573 | -0.834 | 0.404 | 0.202 | 1.907 |
| Sterile-Isolated | 0.204 | 0.792 | -2.005 | **0.045** | 0.043 | 0.965 |
| *High pathogen concentration* (0.1 OD) | | | | | | |
| Amputated-Isolated | 0.707 | 0.460 | -0.754 | 0.451 | 0.287 | 1.742 |
| Infected-Isolated | 2.236 | 0.399 | 2.016 | **0.044** | 1.023 | 4.887 |
| Sterile-Colony | 0.340 | 0.549 | -1.962 | **0.050** | 0.116 | 0.999 |
| Sterile-Isolated | 0.153 | 0.777 | -2.414 | **0.016** | 0.033 | 0.703 |

For the low pathogen concentration, Random factor: Colony, Variance = 0.63, Std. Dev. = 0.80. Likelihood ratio test of model vs intercept-only model: $\chi^2 = 22.30$, df = 5, $p < 0.001$. For the high pathogen concentration, random factor: Colony, Variance = 0.020, Std. Dev. = 0.14. Likelihood ratio test of model vs intercept-only model: $\chi^2 = 28.24$, df = 5, $p < 0.001$. Coefficient estimates are exponentially transformed. P-values below 0.05 are highlighted in bold.

**Table S2.** Statistical differences in ant survival in isolation based on amputation status across different pathogen concentrations. Ants that were infected but not amputated within the colony served as the reference level for both comparisons. We fitted a Mixed Effect Cox proportional hazards model by adding the colony as an intercept random effect.

| Comparison | estimate | std.error | statistic | p-value | conf.low | conf.high |
|---|---|---|---|---|---|---|
| *Low pathogen concentration* (0.01 OD) | | | | | | |
| Infected - Amputated | 0.559 | 0.768 | -0.756 | 0.449 | 0.124 | 2.520 |
| Sterile - Non-amputated | 0.397 | 0.842 | -1.098 | 0.272 | 0.076 | 2.066 |
| Sterile - Amputated | 0.588 | 1.011 | -0.525 | 0.600 | 0.081 | 4.269 |
| *High pathogen concentration* (0.1 OD) | | | | | | |
| Infected - Amputated | 0.134 | 0.731 | -2.751 | **0.006** | 0.032 | 0.561 |
| Sterile - Non-amputated | 0.260 | 1.185 | -1.137 | 0.255 | 0.025 | 2.652 |
| Sterile - Amputated | 0.059 | 0.798 | -3.549 | **<0.001** | 0.012 | 0.281 |

For the low pathogen concentration, Random factor: Colony, Variance = 1.79, Std. Dev. = 1.34. Likelihood ratio test of model vs intercept-only model: $\chi^2 = 10.50$, df = 4, p = 0.03. For the high pathogen concentration, Random factor: Colony, Variance = 0.10, Std. Dev. = 0.32. Likelihood ratio test of model vs intercept-only model: $\chi^2 = 11.95$, df = 4, p = 0.01. Coefficient estimates are exponentially transformed. P-values below 0.05 are highlighted in bold.

**Table S3.** Statistical differences in the survival of ants with amputations at different time points. We fitted a Mixed Effect Cox proportional hazards model by adding the colony as an intercept random effect.

| Comparison | estimate | std.error | statistic | p-value | conf.low | conf.high |
|---|---|---|---|---|---|---|
| Infected vs Sterile | 0.027 | 0.517 | -6.990 | **<0.001** | 0.010 | 0.074 |
| Infected vs Infected+Amputated 0h | 0.370 | 0.357 | -2.787 | **0.005** | 0.184 | 0.744 |
| Infected vs Infected+Amputated 6h | 0.295 | 0.356 | -3.427 | **0.001** | 0.147 | 0.593 |
| Infected vs Infected+Amputated 12h | 0.813 | 0.285 | -0.726 | 0.468 | 0.465 | 1.421 |
| Infected vs Infected+Amputated 24h | 0.676 | 0.303 | -1.293 | 0.196 | 0.373 | 1.224 |

Random factor: Colony, Variance = 0.0002, Std. Dev. = 0.004. Likelihood ratio test of model vs intercept-only model: $\chi^2 = 131.11$, df = 6, $p < 0.001$. Coefficient estimates are exponentially transformed. P-values below 0.05 are highlighted in bold.

**Table S4.** CHC relative abundances (mean ± standard deviation) and their corresponding retention index (RI) for healthy ants and ants with infected and sterile wounds at 2h and 24h.

| | | 0 hours | 2 hours | | | 24 hours | | |
|---|---|---|---|---|---|---|---|---|
| Compound | RI | Healthy | Healthy | Infected | Sterile | Healthy | Infected | Sterile |
| C21 | 2,100 | 0.29±0.14 | 0.31±0.12 | 0.40±0.29 | 0.39±0.17 | 0.32±0.14 | 0.29±0.13 | 0.35±0.08 |
| C22 | 2,200 | 0.24±0.11 | 0.20±0.03 | 0.24±0.12 | 0.22±0.04 | 0.20±0.04 | 0.23±0.07 | 0.20±0.03 |
| C23ene | 2,273 | 0.15±0.05 | 0.17±0.06 | 0.20±0.07 | 0.20±0.09 | 0.13±0.06 | 0.15±0.08 | 0.13±0.02 |
| C23 | 2,300 | 4.96±1.26 | 5.71±1.41 | 7.16±3.40 | 6.12±1.43 | 7.02±0.98 | 6.09±1.37 | 7.40±0.76 |
| 3-MeC23 | 2,373 | 0.14±0.06 | 0.13±0.05 | 0.14±0.05 | 0.16±0.06 | 0.11±0.04 | 0.13±0.05 | 0.11±0.01 |
| C24 | 2,400 | 1.18±0.17 | 1.33±0.17 | 1.44±0.38 | 1.39±0.15 | 1.62±0.14 | 1.52±0.11 | 1.51±0.16 |
| C25diene | 2,470 | 0.10±0.14 | 0.07±0.06 | 0.12±0.08 | 0.11±0.06 | 0.06±0.05 | 0.04±0.06 | 0.11±0.03 |
| C25ene_1 | 2,474 | 1.60±0.75 | 2.03±1.07 | 2.50±1.24 | 2.36±1.39 | 1.96±1.31 | 1.47±1.16 | 2.55±0.79 |
| C25ene_2 | 2,481 | 0.17±0.09 | 0.23±0.10 | 0.26±0.11 | 0.24±0.11 | 0.23±0.09 | 0.21±0.08 | 0.21±0.06 |
| C25 | 2,500 | 11.24±1.90 | 11.57±1.24 | 11.87±1.54 | 11.95±1.60 | 14.57±1.53 | 13.08±1.47 | 13.59±1.18 |
| 13-;11-MeC25 | 2,533 | 0.21±0.09 | 0.14±0.05 | 0.13±0.07 | 0.14±0.06 | 0.10±0.03 | 0.18±0.06 | 0.09±0.03 |
| C26ene; C26diene | 2,575 | 0.52±0.23 | 0.50±0.20 | 0.59±0.22 | 0.57±0.23 | 0.49±0.19 | 0.39±0.16 | 0.61±0.12 |
| C26 | 2,600 | 0.68±0.15 | 0.48±0.10 | 0.51±0.09 | 0.53±0.16 | 0.58±0.13 | 0.61±0.09 | 0.48±0.07 |
| 13-;11-MeC26 | 2,633 | 0.14±0.04 | 0.09±0.03 | 0.07±0.02 | 0.08±0.02 | 0.06±0.02 | 0.09±0.03 | 0.04±0.02 |
| C27ene_1; C27diene | 2,676 | 25.65±3.30 | 28.59±4.65 | 29.44±3.90 | 28.28±5.75 | 27.23±6.23 | 22.30±5.59 | 32.23±4.05 |
| C27ene2 | 2,681 | 0.63±0.59 | 0.46±0.74 | 0.90±1.10 | 0.90±0.65 | 0.34±0.72 | 0.55±0.59 | 0.00±0.00 |
| C27 | 2,700 | 6.57±1.80 | 5.54±1.02 | 5.44±1.09 | 5.78±2.06 | 6.35±1.79 | 6.65±1.48 | 5.33±0.59 |
| 13-; 11-MeC27 | 2,731 | 0.27±0.21 | 0.11±0.04 | 0.13±0.07 | 0.09±0.04 | 0.07±0.05 | 0.10±0.06 | 0.05±0.06 |
| C28ene; C28diene | 2,776 | 1.42±0.60 | 0.95±0.12 | 0.87±0.14 | 0.91±0.13 | 0.79±0.13 | 0.91±0.20 | 0.79±0.10 |
| C28 | 2,800 | 0.69±0.20 | 0.55±0.26 | 0.42±0.14 | 0.53±0.33 | 0.51±0.24 | 0.72±0.28 | 0.36±0.11 |
| 13-; 11-MeC28 | 2,833 | 0.10±0.06 | 0.08±0.06 | 0.06±0.03 | 0.10±0.07 | 0.07±0.05 | 0.08±0.05 | 0.09±0.07 |
| C29ene; C29diene | 2,877 | 27.21±4.10 | 26.57±2.89 | 24.57±3.38 | 25.06±2.28 | 23.23±1.98 | 24.85±2.42 | 22.43±2.05 |
| C29 | 2,900 | 5.49±2.28 | 4.53±2.65 | 3.78±1.73 | 4.43±2.88 | 4.66±2.27 | 6.22±2.04 | 3.04±1.19 |
| 15-; 13-; 11-MeC29 | 2,931 | 0.16±0.22 | 0.01±0.02 | 0.03±0.06 | 0.02±0.04 | 0.00±0.01 | 0.02±0.05 | 0.00±0.00 |
| 3-MeC29; C30ene | 2,976 | 0.82±0.26 | 0.63±0.12 | 0.59±0.13 | 0.66±0.14 | 0.49±0.13 | 0.70±0.19 | 0.47±0.08 |
| C30 | 3,000 | 0.26±0.07 | 0.26±0.16 | 0.23±0.11 | 0.42±0.38 | 0.23±0.11 | 0.31±0.17 | 0.55±0.34 |
| C31ene; C31diene | 3,079 | 7.77±1.63 | 7.54±2.05 | 6.88±1.81 | 7.16±2.22 | 7.48±2.38 | 10.16±2.50 | 6.34±1.70 |
| C31 | 3,100 | 0.76±0.39 | 0.67±0.34 | 0.53±0.21 | 0.67±0.47 | 0.62±0.34 | 1.00±0.34 | 0.49±0.29 |
| C33ene | 3,277 | 0.58±0.08 | 0.55±0.11 | 0.50±0.14 | 0.53±0.16 | 0.49±0.10 | 0.96±0.55 | 0.45±0.14 |

**Table S5.** PERMANOVA analysis comparing CHC composition across wound types and time. The permutations (n = 9999) were not restricted to any group.

| Factor | Degrees of Freedom | Sum of Squares | $R^2$ | F-value | p-value |
|---|---|---|---|---|---|
| Treatment | 2 | 0.046 | 0.07 | 2.702 | **0.035** |
| Time | 2 | 0.055 | 0.08 | 3.252 | **0.017** |
| Treatment x Time | 2 | 0.067 | 0.10 | 3.993 | **0.007** |

P-values below 0.05 are highlighted in bold.

**Table S6.** Pairwise comparisons from PERMANOVA tests for different wound type combinations over time. The false discovery rate was controlled using Benjamini & Hochberg's correction on the computed *p*-values. Permutations (*n* = 9999) were unrestricted across groups. *p.value*: uncorrected *p*-value; *p.adjusted*: false discovery rate-corrected *p*-value.

| Pairwise comparison | $R^2$ | p-value | p-adjusted |
|---|---|---|---|
| Healthy 0h vs Healthy 24 | 0.197 | 0.009 | 0.066 |
| Healthy 0h vs Healthy 2h | 0.060 | 0.324 | 0.377 |
| Healthy 0h vs Infected 24h | 0.143 | 0.039 | 0.118 |
| Healthy 0h vs Infected 2h | 0.144 | 0.037 | 0.118 |
| Healthy 0h vs Sterile 24h | 0.400 | 0.000 | **0.004** |
| Healthy 0h vs Sterile 2h | 0.085 | 0.180 | 0.236 |
| Healthy 24h vs Healthy 2h | 0.119 | 0.101 | 0.151 |
| Healthy 24h vs Infected 24h | 0.156 | 0.062 | 0.120 |
| Healthy 24h vs Infected 2h | 0.092 | 0.148 | 0.208 |
| Healthy 24h vs Sterile 24h | 0.145 | 0.072 | 0.124 |
| Healthy 24h vs Sterile 2h | 0.074 | 0.234 | 0.289 |
| Healthy 2h vs Infected 24h | 0.169 | 0.050 | 0.120 |
| Healthy 2h vs Infected 2h | 0.040 | 0.447 | 0.494 |
| Healthy 2h vs Sterile 24h | 0.237 | 0.020 | 0.085 |
| Healthy 2h vs Sterile 2h | 0.015 | 0.774 | 0.795 |
| Infected 24h vs Infected 2h | 0.239 | 0.013 | 0.066 |
| Infected 24h vs Sterile 24h | 0.429 | 0.002 | **0.016** |
| Infected 24h vs Sterile 2h | 0.148 | 0.063 | 0.120 |
| Infected 2h vs Sterile 24h | 0.125 | 0.077 | 0.124 |
| Infected 2h vs Sterile 2h | 0.019 | 0.795 | 0.795 |
| Sterile 24h vs Sterile 2h | 0.153 | 0.054 | 0.120 |

P-values below 0.05 are highlighted in bold.
